# Cardiac Spiral Wave Drifting Due to Spatial Temperature Gradients – a Numerical Study

**DOI:** 10.1101/362913

**Authors:** Guy Malki, Sharon Zlochiver

**Affiliations:** Department of Biomedical Engineering Department, Faculty of Engineering, Tel-Aviv University, Ramat-Aviv, Tel-Aviv, 69978, Israel

**Keywords:** AF model, Computer simulations, Guided atrial ablation, Rotor meandering and drifting

## Abstract

Cardiac rotors are believed to be a major driver source of persistent atrial fibrillation (AF), and their spatiotemporal characterization is essential for successful ablation procedures. However, electrograms guided ablation have not been proven to have benefit over empirical ablation thus far, and there is a strong need of improving the localization of cardiac arrhythmogenic targets for ablation. A new approach for characterize rotors is proposed that is based on induced spatial temperature gradients (STGs), and investigated by theoretical study using numerical simulations. We hypothesize that such gradients will cause rotor drifting due to induced spatial heterogeneity in excitability, so that rotors could be driven towards the ablating probe. Numerical simulations were conducted in single cell and 2D atrial models using AF remodeled kinetics. STGs were applied either linearly on the entire tissue or as a small local perturbation, and the major ion channel rate constants were adjusted following Arrhenius equation. In the AF-remodeled single cell, recovery time increased exponentially with decreasing temperatures, despite the marginal effect of temperature on the action potential duration. In 2D models, spiral waves drifted with drifting velocity components affected by both temperature gradient direction and the spiral wave rotation direction. Overall, spiral waves drifted towards the colder tissue region associated with global minimum of excitability. A local perturbation with a temperature of T=28°C was found optimal for spiral wave attraction for the studied conditions. This work provides a preliminary proof-of-concept for a potential prospective technique for rotor attraction. We envision that the insights from this study will be utilize in the future in the design of a new methodology for AF characterization and termination during ablation procedures.

## 1. INTRODUCTION

Atrial fibrillation (AF) is the most common sustained cardiac arrhythmia, affecting more than 10% of the elderly population [1]. AF is characterized by rapid and irregular activation of the atrium, and is often the result of fibrillatory conduction maintained by the existence of one or few organized “mother rotors”, or alternatively by the existence of focal ectopic sources, [2-3]. Pharmacological treatment for AF includes antiarrhythmic drugs that are either rate or rhythm control. These effects are achieved by various ionic mechanisms altering the electrophysiological properties of the membrane voltage at either depolarization or repolarization phases, e.g. by reducing cellular excitability, or by increasing the refractory period. Nonetheless, as pharmacological treatment for AF shows limited success [4] and due to its possible long term negative side-effects, surgical ablation procedures have become attractive for curing AF in symptomatic patients. These procedures try to isolate and annihilate arrhythmogenic sources in the atrial tissue by inducing permanent tissue ablation, typically by delivering radiofrequency (RF) energy. While unguided, empirical ablation is practiced in some procedures, it often results in redundant applications of RF energy that may cause inadvertent injury and thromboembolic complications in the tissue. Thus, the correct characterization of the arrhythmogenic sources (rotors vs. focal activity) and their localization are of the utmost importance for a successful outcome of the ablation procedure, and several electrogram-guided ablation techniques have been recently proposed or utilized for that purpose [5-6]. Those techniques are based on either time-domain or frequency-domain analysis, and constructed by single electrogram analysis, or multiple, normally simultaneously-recorded electrograms, to provide spatiotemporal characterization of the electrical activity in the atrial tissue. Electrogram-derived indices, such as complex fractionated atrial electrograms (CFAEs) [7-8] and dominant frequency (DF) [9-10], have been proposed as guide to perform electrogram-based ablation, and recently logical combination of rate and regularity measures has been developed to achieve computational mapping of the atrial sources [6]. Phase analysis using phase singularity evaluation has been used to characterize rotors and their pivot points [5, 11].

Though, none of these practices were successfully implemented in clinical settings because of various limitations in their ability to accurately characterize the arrhythmogenic source zones due to noise, misleading phase and activation times that distort the reconstructed maps [11]. Moreover, novel techniques that involve more advanced signal processing methodologies to locate the pivot points of persistent rotors were proposed lately, including principal component analysis [12], and spatial Shannon entropy measurement [13]. Nevertheless, due to the yet poor understanding and the ambiguity of the correlations between the underlying arrhythmogenic activity and its electrogram manifestation, reported ablation success rates are similar for electrogram-guided as for empirical ablation procedures [4, 14]. Therefore, there is a clear need for new approaches to improve guided ablation procedures by better detection and characterization of arrhythmogenic drivers in order to minimize the destruction of unnecessary healthy tissue and allow for higher long-term success rates of such procedures.

Here we propose a novel strategy for addressing this need for the specific cardiac rotors arrhythmogenic drivers in persistent AF patients, which is based on artificial induction of temperature gradients in the atrial tissue. The effect of temperature on the biophysical properties of various biological tissues has been long studied. Temperature sustains direct impact on biological processes, thus influencing ion channel kinetics, action potential (AP) morphology and other electrophysiological properties via its effects on the rate constants of chemical reactions and sub-cellular biological processes [15-17]. These effects are commonly modelled by the multiplication of the relevant reaction rate constants (e.g., ion channel gating variables) by a scaling factor, which sustains a power-law relationship with temperature following Arrhenius’ law [18]. Consequently, e.g., a temperature increase results in faster gating kinetics leading to a decrease in the cardiac AP duration (APD), while induced hypothermia on the other hand inhibits the dynamics of the membrane currents, effectively prolonging the APD [19]. Several studies have related temperature changes to cardiac pathologies associated with impaired electrical conduction [20-23]. The contribution of temperature on pro-arrhythmic APD restitution properties and alternans formation was recently studied by Fenton et al. [24-25] using simplified ventricular model with thermoelectric coupling. Their studies showed that alternans and conduction block onset occur in higher cycle lengths as temperature decreases, and that the minimum tissue size required to sustain hypothermic ventricular fibrillation is temperature dependent. Nevertheless, ionic mechanisms explaining the effect of temperature could not be pursued due to the abstract modelling approach. Finally, Yamazaki et al. [26] demonstrated in an experimental study using rabbit hearts that regional cooling within the left ventricle facilitates termination due to collisions with boundaries of spiral-wave re-entry through unpinning of rotors and drift toward the cooled region. Here too, these important findings lack detailed biophysical and theoretical elucidation, and are supported by experimental evidence only. In this study we will examine a possible positive effect of externally applied spatial temperature gradients (STGs) on atrial tissues activated by a dominant rotor – induced controlled drifting. While the exact mechanism remains unclear, both experimental and numerical studies show rotor drifting and meandering due to various types of spatial gradients in both biophysical and anatomical properties [27-29]. Rotors have been shown to drift towards regions of lower excitability of various origins, including heterogeneity in sodium channel availability [30-31], or in IK1 channel density [32]. Based on these results, we postulate that externally applied STGs will enable controlled rotor drifting by a similar mechanism of spatial heterogeneity in atrial excitability. The aim of this study is to establish the theory underlying the relationship between temperature gradients, rotor drifting and tissue excitability. We envision that in the future our approach can indeed promote relevant clinical applications during ablation procedures.

## 2. METHODS

### A. Biophysical Modeling

Atrial electrical activity was simulated in either a single cell or a 30 mm X 30 mm 2D tissue by solving the following reaction-diffusion equation by adopting the mono-domain formalism and under the approximation of tissue isotropy:

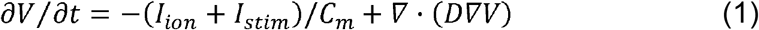

where *V* [mV] ¡s the transmembrane voltage, *C*_*m*_ [μF/cm^2^] is the membrane capacitance per unit area, *I*_*stim*_ and *I*_*ion*_ [μA/cm^2^] are the external stimulation and membrane ionic currents, respectively, and *D* [mm^2^/ms] is the diffusion coefficient. Human atrial kinetics were employed for calculating *I*_*ion*_ using the Courtemanche-Ramirez-Nattel (CRN) model [33]. In most simulations, the CRN model was modified to account for chronic AF remodeling by downregulating the channel densities of the transient outward K+ current (*I*_*to*_) by 50%, the ultra-rapid delayed rectifier K+ current (*I*_*Kur*_) by 50%, the L-type Ca2+ current (*I*_*Ca–L*_) by 70%, and by increasing the density of the inward rectifier K+ current (*I*_*K1*_) by 100%, according to previous publications [30, 34]. The diffusion coefficient was adjusted to *D* = 0.03 mm^2^/ms to achieve planar conduction velocity (CV) of ~0.4 m/s. The effect of temperature on the model kinetics was incorporated by multiplying the rate constants *α*(37°*C*) and *β*(37°*C*) of the gating variables of several ion channels at a reference temperature of 37°*C* by the temperature adjustment factor *Q* (*T*) following Arrhenius equation, as originally proposed by Hodgkin & Katz [15]:

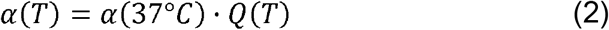

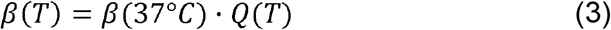

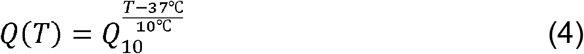

where *T* [°*C*] is the local tissue temperature, and *Q*_10_ is the temperature coefficient defining the ratio by which the rate constants increase following a temperature increase of 10°C. Equations (2-4) were applied to the six main ion channel currents as detailed in Table 1. Equation (1) was solved numerically by employing forward Euler integration in time and the finite difference method in space, using temporal and spatial resolutions of Δ*t* = 2.5 and Δ*h* = 0.1 mm, respectively. Those values were set after ensuring convergence of the simulation data by comparison to those obtained with finer resolution values

**Table 1.** Qio VALUES FOR EACH IONIC CURRENT

| Current | $Q_{10}$ | Reference |
| --- | --- | --- |
| $I_{Na}$ | 3 | [33] |
| $I_{to}$ | 2.2 | [34] |
| $I_{Kur}$ | 2.2 | [34] |
| $I_{Kr}$ | 3.3 | [35] |
| $I_{KS}$ | 2.57 | [36] |
| $I_{Ca-L}$ | 2.2 | [37] |

### B. Simulation Configuration

Single cell simulations were conducted to study the effect of temperature on basic electrophysiological properes. Spiral waves were established in a uniform 2D geometry with a fixed temperature of 37°C, using the S1-S2 cross-field stimulation. This protocol induced a counter clockwise or clockwise rotating spiral wave in the middle of the tissue with a dominant activation frequency of 8.1 Hz. Five second long electrical activity was simulated prior to any application of spatial temperature gradients (STGs) to ensure initial stability of the spiral wave. At the end of that stabilization period, STGs of either one of two types were applied (see Fig. 1): 1) linearly changing temperature gradients along the y-axis between *T*_*1*_ = 37ΰC and varying *T*_2_ (between 30°C and 44°), so that and *T*(*y* = 0) = 37°C and *T*(*Y* = 30*mm*) = *T*_2_.

**Fig. 1.**
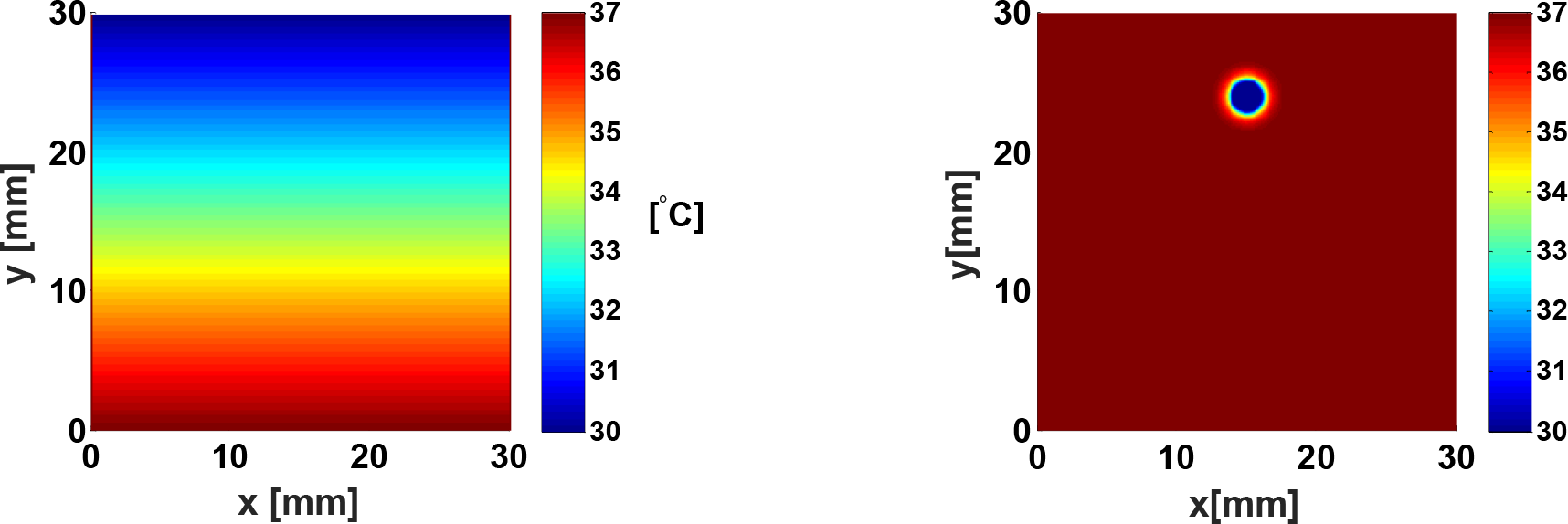
Two types of spatial temperature gradients that were employed in the study. Left –a linear temperature gradient, in this example with *T*_2_ = 30°C, right –a local circular temperature perturbation with a radius of 1mm, here with *T*_2_ = 30°C

This configuration allowed the investigation of basic spiral wave drifting mechanisms under controlled temperature gradients; 2) local circular temperature perturbation with a radius of 1mm, typical for AF ablation catheters. This configuration was intended to mimic the potential clinical application of an external heat source catheter during an EPS procedure. Two types of perturbation configurations were simulated: in the first type, the location of the perturbation center was kept fixed at 8mm from the initial location of the spiral wave core center, but its temperature was varied between 20°C and 36°C. In the second type, the perturbation temperature was fixed at 28, but its distance from the core of the spiral wave was changed between 5mm and 11mm. To account for the heat distribution due to the application of a circular heat perturbation, the following bio-heat transfer equation was numerically solved in steady-state [40]:

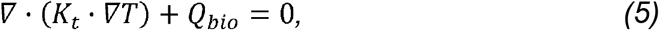

where *K*_*t*_ = 0.7 W/(m·K) is the atrial tissue thermal conductivity, and *Q*_*bi0*_ [W/m^3^] is a source term representing the blood perfusion, and is equal to:

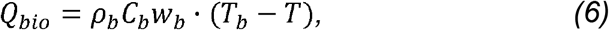

where *ρ*_*b*_ = 1080 kg/m^3^ is the blood density, *C*_*b*_ = 3600 J/(kg·K) is the specific heat capacity of the blood, *w*_*b*_ = 1 Hz is the perfusion rate and *T*_*b*_ = 37°C is the arterial blood temperature. Equations (5-6) were solved numerically for each perturbation configuration in Fig. 1 using COMSOL finite element software (COMSOL, Burlington, MA).

### C. Technical Aspects and Data Analysis

All simulations were performed using C++ code on a high-performance cluster computer (Altix X86-PTO; Silicon Graphics International, Milpitas, CA) with a master node (eight cores, Xeon 2.5 GHz processor; Intel, Santa Clara, CA) and up to five computational nodes (60cores, Xeon 2.8 GHz; Intel). Data analysis and visualization were performed with MATLAB R2014b (The MathWorks, Natick, MA). In order to investigate the spiral wave stability and its rate of drifting, tip trajectory was automatically tracked following Nayak et al. [41]. Briefly, this method locates the singularity point as the intersection region between a certain iso-potential range (in our case between −40mV and −15mV, range A) and the range *ϑV/ϑt* = 0 (range B). To avoid incorrectly detecting overlapping points, we first employed a 3×3 pixel median filter over the range A to fill in small gaps, and then defined the singularity point map as the overlapping areas of the filtered range A and range B (range C). Since in most cases more than one potential singular point was detected in range C, we used the exponentially-weighted moving average (EWMA) method to choose the most likely singular point in that specific time frame. In the EWMA method, we first set exponentially decreasing weights to all singular points detected in previous time frames, with the highest weight given to the most recent detected singular point. We then calculated a projected location of the next singularity point by employing a moving average on the exponentially weighted past singularity point locations. Lastly, we selected the singularity point out of range C that was the closest to the projected location. Spatial excitability distribution was represented by the sodium channel availability, given by the product of its fast (*h*) and slow (*j*) inactivation gates [2].

## 3. RESULTS

### Single Cell Simulations and Model Validation

The effects of temperature on the AP morphology and on cellular excitability are shown in Fig. 2, for both standard CRN model parameters (panel A) and for chronic AF modifications (panel B). Temperature variations sustained significant effects in standard model parameters, as can be seen from a detailed characterization of the temperature dependence in the ionic model in panel C, which presenting the analysis of these parameters at a vast range of temperatures that does not produce irreversible tissue damage (20°C-42°C). Both APD at 90% repolarization (APD90), as well as recovery time from inactivation, calculated by the time interval between sodium channel availability (*h*×*j*) levels of 0.1 and 0.9, (panel C, upper and lower left columns, respectively), decreased with increasing temperatures. These results are a direct consequence of the increasing rate constants with increased temperatures (2-3), and support our hypothesis assumption that low temperature regions in the tissue exhibit reduced excitability. When chronic AF modifications were implemented, the effect of temperature variations on the AP morphology and duration was minor. Notwithstanding, temperature effect on the recovery time from inactivation was still substantial, signifying that the correlation between low temperature and reduced excitability holds in chronic AF conditions as well. The AP amplitude as function of temperature (panel C, upper right plot) exhibits a similar behavior, in which decreased as the temperature increasing. This observation attests that the sodium channel rate constant effected the sodium gating to open more slowly in a colder temperature, and therefore amplified the depolarization process. On the other hand, CV (panel C, bottom right plot) raised as the temperature increased, because of the growing rate constants, which yielded to rapider propagation. Furthermore, spiral wave dynamics with constant tissue temperature have been explored, as can be seen in Fig. S1. Three spiral wave properties were analyzed to describe the tip meandering as function of temperature: the fast frequency that determines the interval between consecutive local excitation, also called the activation frequency (*f*_1_); the slow frequency that determines the rotation rate of the spiral wave tip, denoted as the meander frequency (*f*_2_); and the mean tip velocity during single and complete rotation cycle. All of them are aligned with the model predications, so that the spiral wave dynamics are reduced, became slower and have lower frequencies, as the tissue temperature is decreasing.

**Fig. 2.**
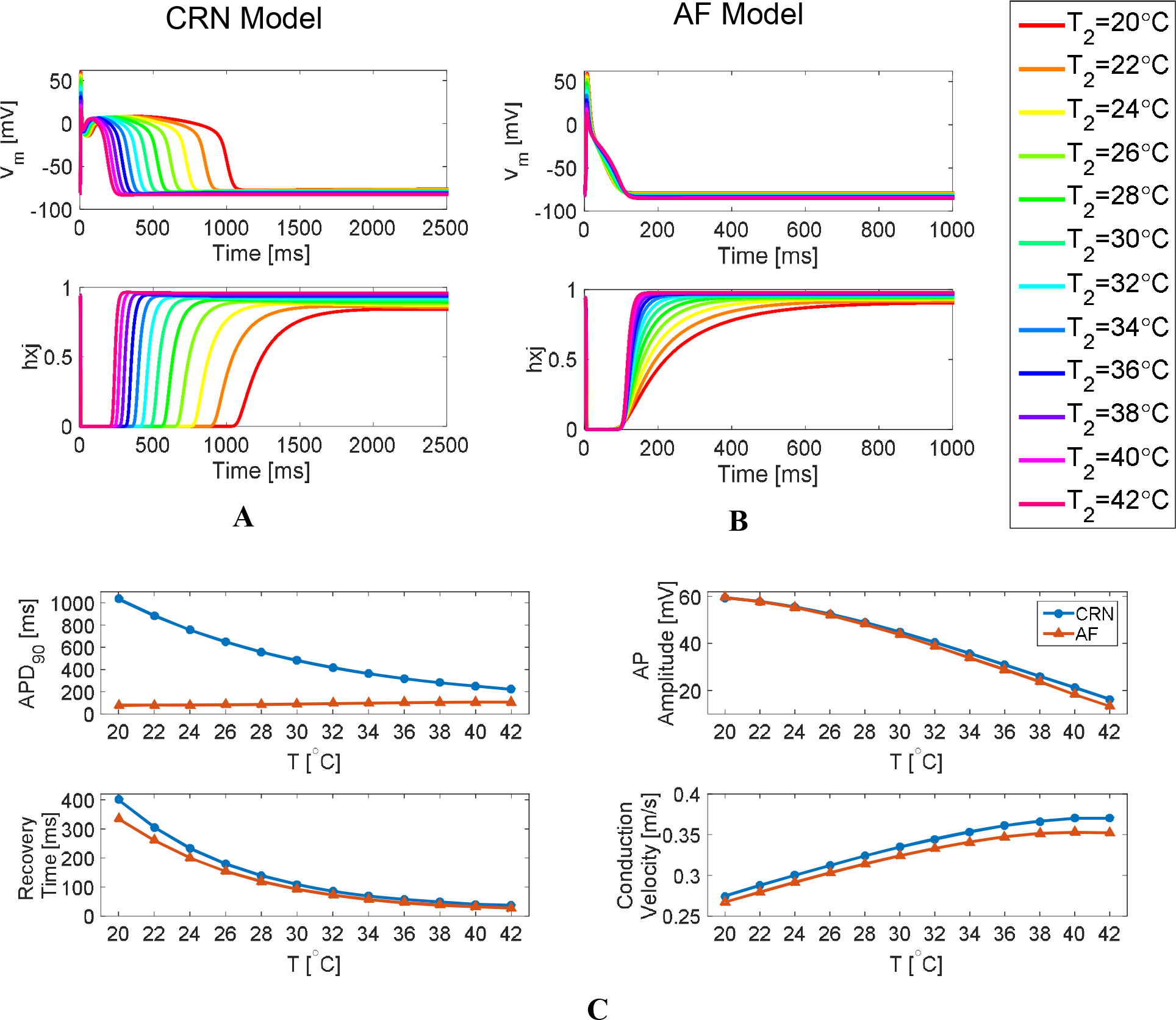
Effects of temperature dependency on single cell action potential and excitability. A-B. Transmembrane voltage (top) and sodium channel availability (h×j) in time for various temperatures and for the standard CRN model (A, [33]) and the AF model (B, [34]). C. Temperature dependency on the model outcomes for the two kinetic models: APD_90_ (top left), sodium channel recovery time (bottom left) measured as the time interval between hxj of 0.1 and 0.9, AP amplitude (top right), and planar conduction velocity (bottom right).

### Tissue Simulations – Linear Temperature Gradients

2D simulations were conducted for a 30 mm X 30 mm atrial tissue model using the CRN model with chronic AF modifications. Vertical linear STGs were applied as detailed in the methods section. Fig. 3A shows typical tip trajectories due to either a positive STG (*T*_2_ = 40°c > 37°C, in green) or a negative STG (*T*_2_ = 34°C < 37°C, in blue), for both clockwise and counterclockwise rotation (light and dark traces, respectively). It can be seen that the vesical drifting component was directed opposite to the gradient direction (*dT/dy*), while the horizontal component was directed according to the sign of the product (*dT/dy*) · *R*, where *R* obtains the values of +1 or −1 for clockwise and counter-clockwise rotations, respectively. For example, a clockwise rotating spiral wave (*R* = 1) with *T*_2_ = 34°C (*dT/dy* < 0) drifted towards the upper-left corner (light blue trace) of the tissue due to a positive vertical component (– *dT/dy* > 0) and a negative horizontal component (*dT/dy* · *R* < 0). Fig. 3B illustrates the spatiotemporal tip trajectory for a clockwise rotating spiral wave and for linear STGs different by their *T*_*2*_ varying between 30°C and 44°C. Mean tip trajectories (bold lines) were calculated by applying spatial trajectory averaging using sliding temporal windows of durations that corresponded with the time required by the tip to complete a single rosettelike cycle. The mean vertical and horizontal drifting velocities were calculated from the mean tip trajectories by extracting the total *y*-axis and *x*-axis tip translation during the first 3 seconds of activity, and are shown as a function of (*dT/dy*) in Fig. 3C. Fig. 3D represents the angle between the drift velocity vector and the gradient vector as a function of (*dT/dy*). This analysis reveals the effect of temperature on the drifting angle, and discovers a discontinuity at zero gradient that was emphasized by studying smaller gradients in both directions. It should be noted that at larger gradients (e.g. T2=30°C and T2=44°C), in addition to the change in the drifting angle, the drifting trajectory pattern is slightly altered after the initial drift due to interaction with the boundaries. Thus, Figs. 3B-D demonstrate that wave drifting translation monotonically and symmetrically depended on the difference *T*_2_ – 37°C.

**Fig. 3.**
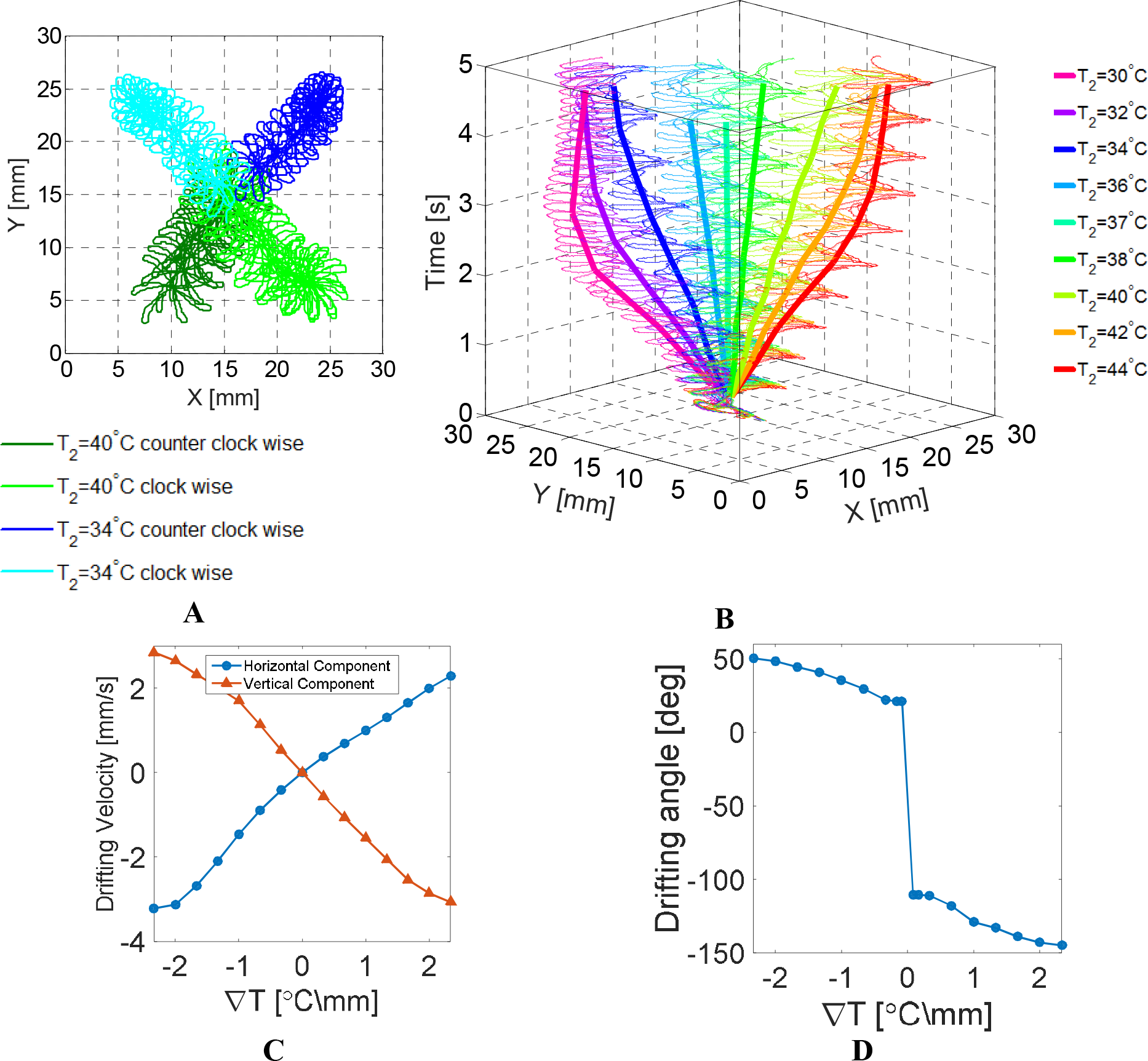
Linear temperature gradients in 2D tissue simulations. A. Effects of temperature gradient sign and direction of spiral wave rotation. Negative and positive gradients are marked in blue and green, respectively. Clockwise and counter clockwise rotations are marked in light and dark shades, respectively. B. Spatiotemporal spiral wave tip trajectory and its mean (light and bold lines, respectively) for linear temperature gradients of various temperatures, T2. All spiral waves rotated clockwise. C. Mean horizontal and vertical drifting velocities as a function of the temperature gradient. D. The angle between the drift velocity vector and the gradient vector as a function of the temperature gradient.

Next, we demonstrate that, in agreement with our hypothesis, spiral wave drifting is due to the induction of spatial heterogeneity in tissue excitability following the application of the STG, such that drifting occurs from high to low excitability regions. We focus on the case in which an STG with T_2_=30°C is applied on a tissue with a clockwise rotating rotor at t=0. As was shown in Fig. 3, the application of such STG resulted in a spiral wave diagonally drifting towards the top-left corner of the tissue. The mean drifting trajectory is shown in Fig. 4A, while the temporal evolution of the sodium channel availability (represented by *h*×*j*) is shown as a time-space plot in Fig. 4B along the diagonal profile that matches the drifting trajectory (dashed line in Fig. 4A). In this plot, the *x*-axis represents a normalized linear coordinate along the diagonal profile (i.e., *p*=0 and *p*=1 represent the top-left and bottom-right edges, respectively), the *y*-axis represents time, and the *h*×*j* values are coded by color. The locations of the spiral wave pivot along the diagonal profile can be easily visualized on the time-space plot as the thick, purple-colored line (marked with an arrow), thus representing low sodium channel availability, that starts close to the center (p=0.5) at t=0, and ends at the top-left region of the tissue (p=0.16) where the drifting ends after ~3 sec. Figure 4C presents a zoomed-in view of part of the time-space plot, and demonstrates the consistent imbalance in the *h*×*j* values between the two sides of the pivot location during the drifting process. For each rotation cycle of the spiral wave the sodium channel availability (hence excitability) to the left of the pivot is lower than to its right (see markers on the inset in Fig. 4C). Hence the pivot movement to the left indicates drifting of the spiral wave towards the region with reduced excitability, i.e., the colder region. To further demonstrate this idea, the 5-sec long time average of the *h*×*j* values along the diagonal profile is shown in Fig. 4D, with the beginning and ending pivot points marked by dots. This figure clearly shows the pivot drifting towards the low excitability region, or more precisely, to the location with globally minimum mean *h*×*j*. To understand the dynamical mechanism by which drifting was occurring towards the low excitability (colder) region of the tissue, an activation map corresponding to one spiral wave rotation during drifting is shown in Fig. 5. This map shows wavefront activation isochronal lines with temporal intervals of 10ms. The tip trajectory during that cycle is marked by the bold black line, with the orange and green dots showing the tip initial and final locations, respectively. The activation map clearly shows the net diagonal drifting towards the top-left corner during that cycle. During drifting from a hotter to a colder region, the isochrones become denser and denser, indicating slowing down of the wavefront due to propagation in a gradually less excitable region. This also implies a longer drifting distance that was required for the wavefront before being able to rotate back towards the hotter region, due to its reduced capability to reenter in a low excitability region in comparison with its increased capability to reenter in a high excitability region. The gradually reducing tissue excitability during the spiral wave drifting towards the colder region was a direct consequence of the longer recovery time associated with the reduced local temperature (see Fig. 2C). Since the imposed STG was linear, and as recovery time increases non-linearly but rather exponentially with decreasing temperatures (Fig. 2C), this also contributed to a longer required tip drifting distance when traveling towards a colder region in comparison to travelling towards a hotter region before the local tissue was sufficiently recovered to sustain a successful reentry. Hence, overall net drifting was towards the colder region.

**Fig. 4.**
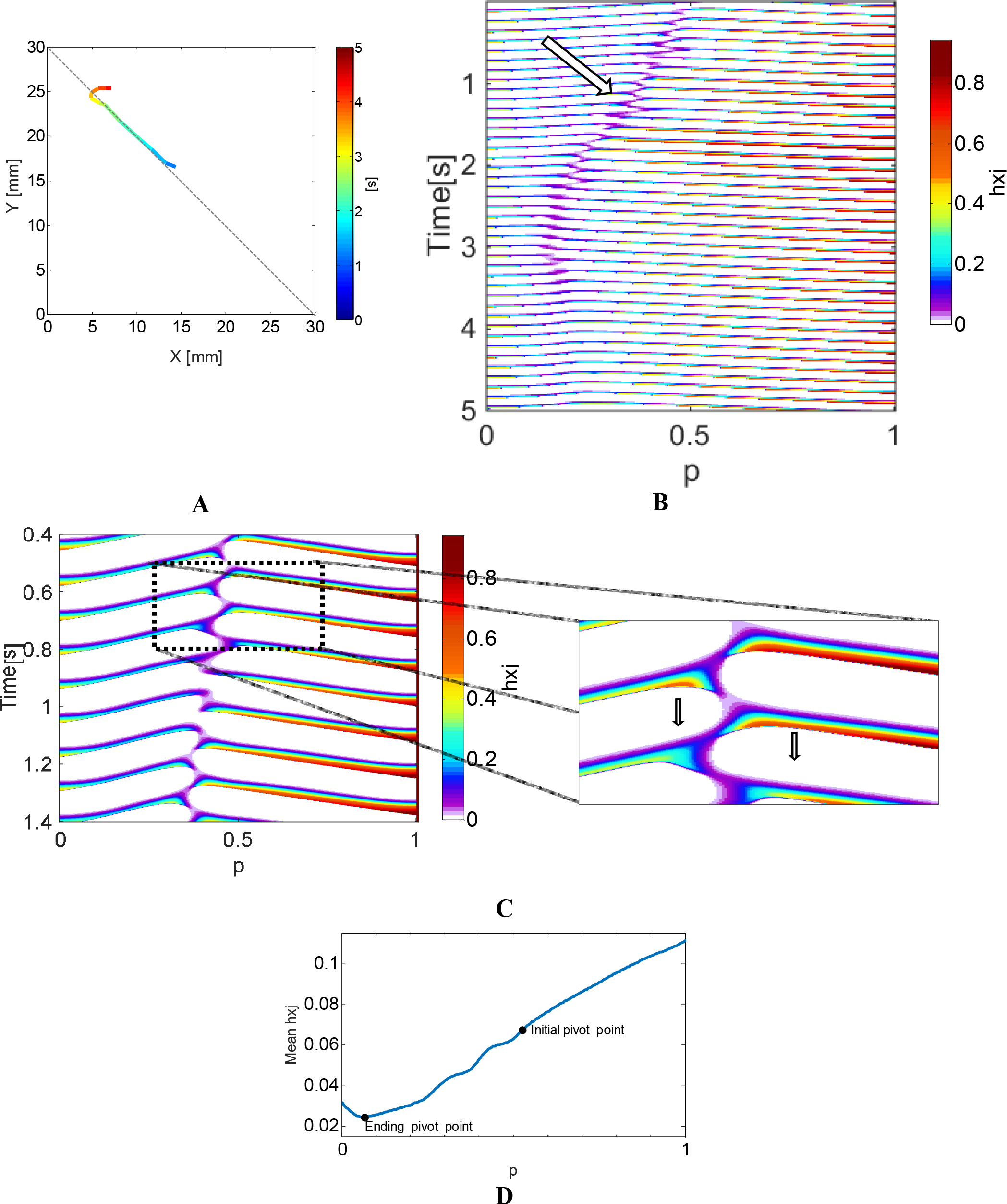
Drifting occurs from high to low tissue excitability in a tissue with a linear temperature gradient. A. Mean spiral wave tip drifting trajectory for a model with a linear temperature gradient with T2=30°C and a clockwise rotating spiral wave. B-C. A time-space plot (TSP) of the sodium channel availability (h×j) along the dashed profile in panel A. The x-axis is represented by a normalized linear coordinate along the diagonal profile (p). A zoomed in section is given in panel C, demonstrating the average lower sodium channel availability to the top-left of the spiral wave core. D. Time-averaged sodium channel availability along the profile marked in panel A, showing that drifting indeed occurred towards the globally minimal average h×j.

**Fig. 5.**
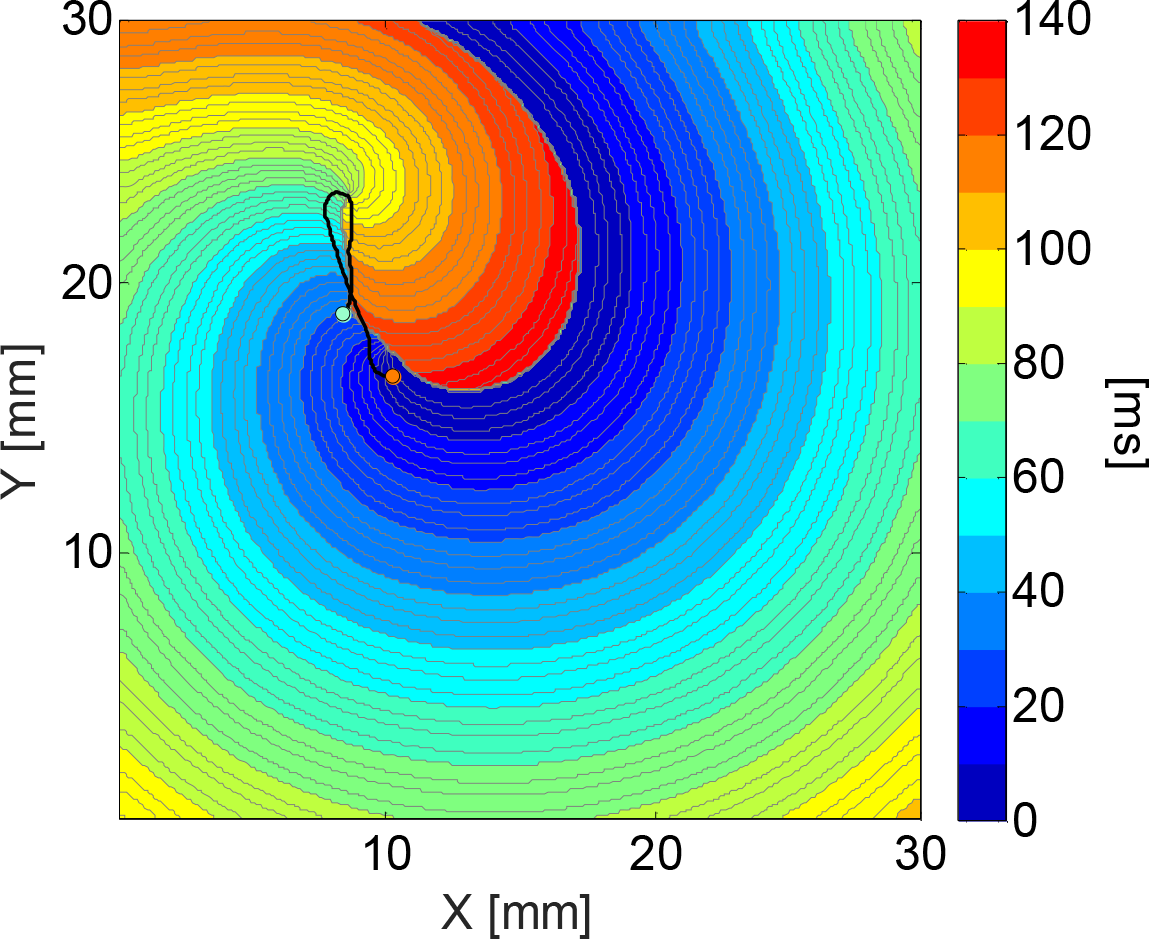
Activation map of a single spiral wave rotation corresponding to the linear temperature gradient simulation with T2=30°C. The orange and green points correspond to the initial and end points of spiral wave tip during this single cycle. A net drift towards the top-left corner is seen during the rotation. See more details in the text.

### Tissue Simulations – Local Perturbations of Varying Temperatures

Next we studied the feasibility of a clockwise rotor attraction by a small temperature perturbation, that may represent, e.g., the application of an external cooling probe. Circular temperature perturbations with a radius of 1mm were modeled as described in the methods section, with a constant atrial background temperature of T_1_=37. Following the results in the previous section, only perturbations having temperature T_2_<T_1_ were modeled since spiral waves drift from high to low temperature regions. Fig. 6A shows the mean tip trajectories during 50 seconds of activity when applying perturbations with various temperatures between 20°C and 36°C, and for the perturbations location kept fixed at a distance of 8mm from the initial location of the spiral wave core center. With the exception of perturbation temperatures of 20°C and 36°C, for which no attraction occurred, rotors were attracted to the cooler perturbation following a three-phase pattern: 1) a slow transient phase in which rotors were first slowly drifting, 2) a fast transient phase in which the rotors were rapidly drifting towards the perturbation, and 3) a steady state phase where the rotors anchored around the perturbation. For perturbation temperatures of 28°C or higher, the slow transient phase 1 was relatively short, and its duration was found to decrease as the perturbation temperature decreased. An example is given in Fig. 6B for T_2_=28°C (the results for all other temperatures are given in Fig. S2). The left panel shows the tip trajectory (in green) and the mean trajectory (in black), while the right panel shows the distance (and mean distance, in black) of the spiral wave tip trajectory from the center of perturbation that is located at (*x*,*y*)=(15, 23mm) as a function of the first 20 seconds of activity, with the three characteristic phases marked. For temperatures lower than 28°C, the slow transient phase was long, characterized by a first detraction and then attraction to the perturbation. Moreover, in contrast to the higher temperatures, the duration of the transient phase increased as the perturbation temperature decreased. An example for this pattern is given in Fig. 6C for the case of T_2_=26°C, Similarly to Fig. 6B. For T2=36°C and T2=20°C no attraction for the RTP occurred. At the T2=36°C case the spiral wave diverted awhile, and returned to the middle of the tissue, without attracting the RTP, due to the small gradient in the heterogeneity of excitability. On the other hand, the rotor trajectory at the T2=20°C case is having a similar pattern to the 22°C-26°C cases, which include a long slow transient. However, for the T2=20°C no attraction established in the time frame of the simulation (50sec).

**Fig. 6.**
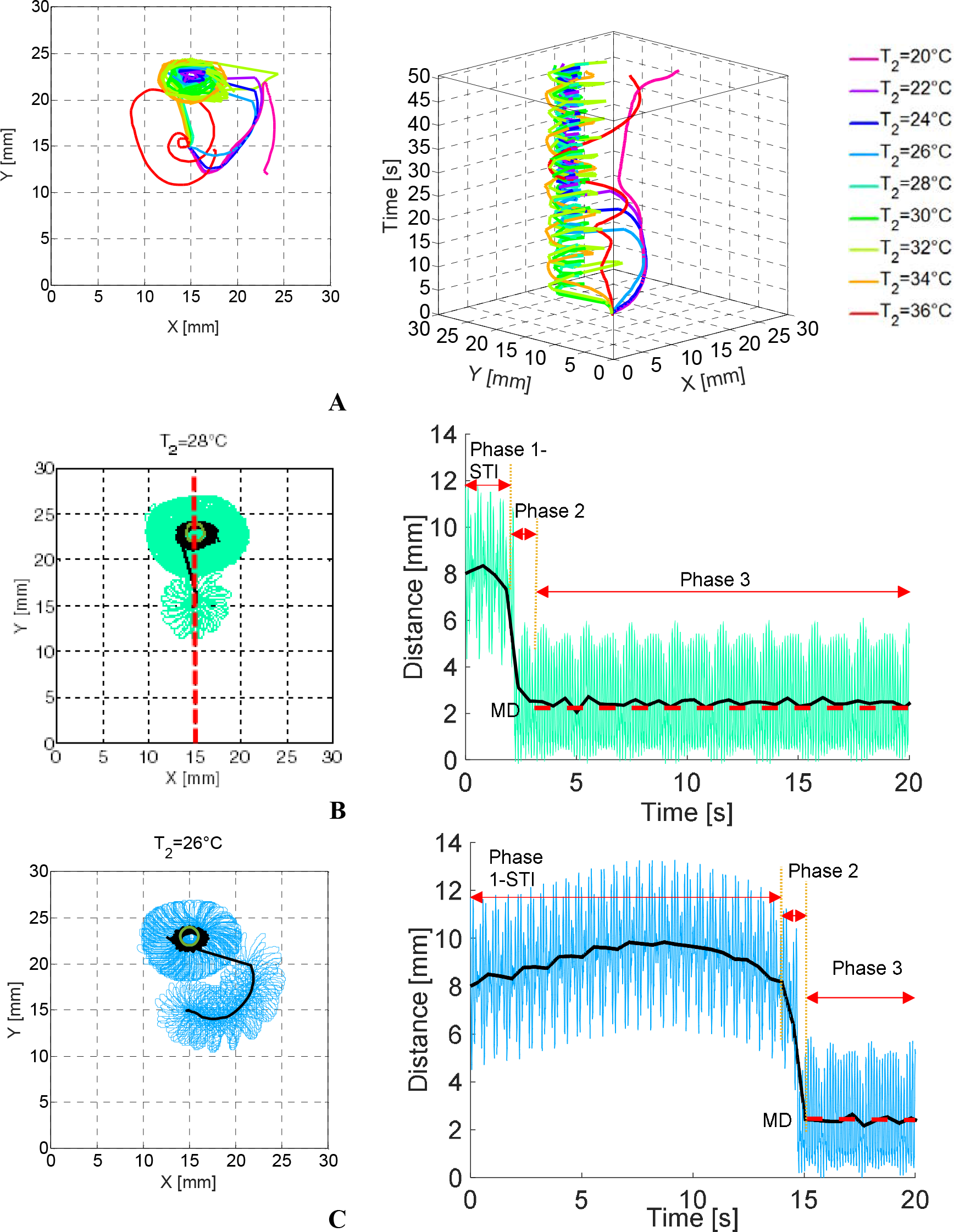
Simulations of local perturbations of varying temperatures. A. Left - mean tip trajectories of a spiral wave in the presence of the local perturbations. Right – time-space plots of the trajectories. B-C. Detailed tip trajectories and mean trajectories (in black) for the cases of T2=28°C and T2=26°C (B and C, respectively). Left – the distance of the mean tip trajectory from the center of perturbation as a function of time. These traces show the typical 3 phases of drifting – slow transient phase (Phase 1), fast transient phase (Phase 2) and steady state phase (Phase 3). The measures of slow transient interval (STI) and mean distance at steady state (MD) are marked.

To better quantify the quality of rotor attraction to the local temperature perturbation, an attraction cost function, *ϕ*, was established that considers both speed of attraction and its steady state properties. Referring to the right panel of Fig. 6B-C we defined *STI* [ms] as the slow transient interval, and *MD* [mm] as the mean distance of the spiral wave from the perturbation center at steady state. To calculate the *STI*, the time derivative of the mean tip distance (i.e., the gradient of the black curve in Fig. 6B, right) was first calculated, and the *STI* was defined as the time corresponding to the maximal derivative. These parameters were calculated for all perturbation temperatures, and in order to account for the mixed units, each was normalized to obtain a mean of 10 with a standard deviation of 3, resulting in corresponding 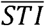 and 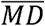. The cost function was calculated as the following root-mean-square:

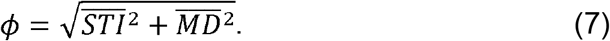

Figure 7A shows *SIT* and *MD* as a function of the perturbation temperature, while panel B shows the cost function values, clearly demonstrating an optimal perturbation temperature of 28°C. At that temperature the optimal balance between fast drifting time to the perturbation and anchoring at a low distance from it was reached.

**Fig. 7.**
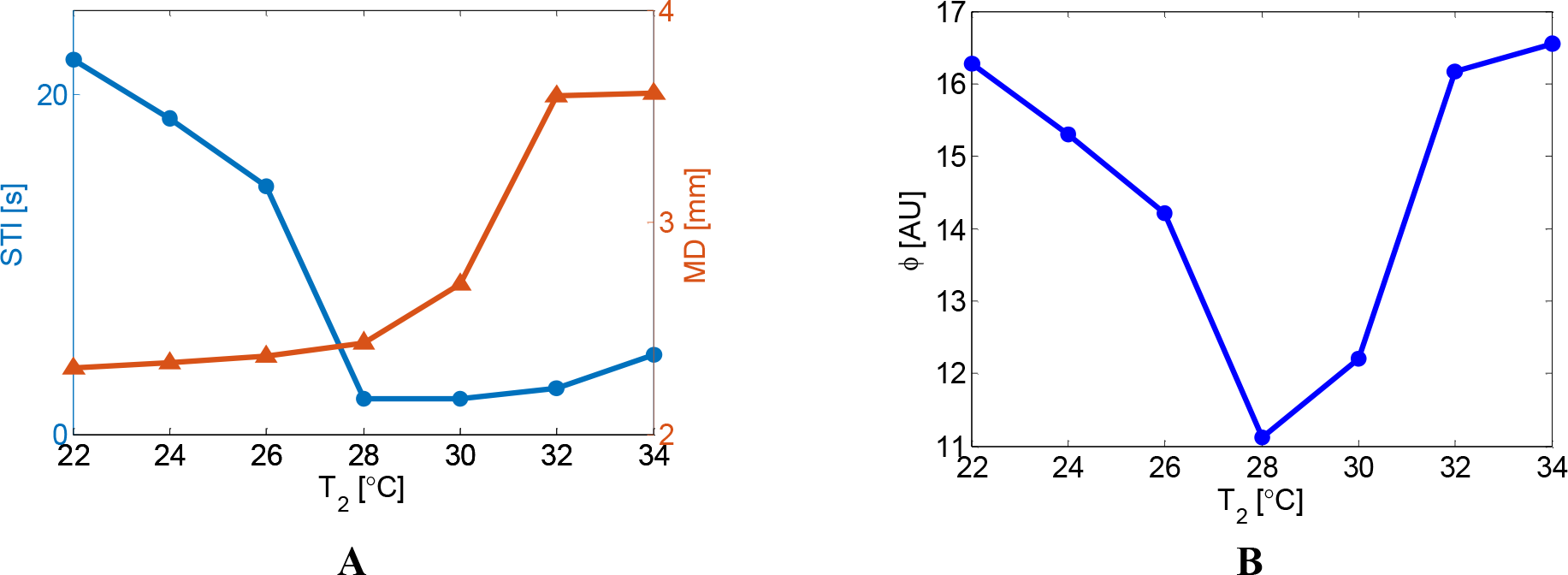
Quality of spiral wave attraction as a function of perturbation temperature. A. Slow transient interval (STI) and mean distance (MD) of the spiral tip. B. Cost function, as defined in Eq. 7, as a function of temperature, showing a clear optimal temperature of 28°C.

In order to check if the optimal temperature changes for perturbations at other pre-set locations, we conducted simulations with varying temperature perturbations using additional distances of 6, 7 and 9mm (which was the maximal distance that enabled attraction at 28°C). The optimal temperature of 28°C was obtained in all these pre-set distances, similarly to the 8-mm case. Hence, we can infer that the optimal temperature that yields fast and stable drifting is 28°C regardless of the pre-set distance of the perturbation from the rotor, and therefore was used in the rest of our simulation study.

### Tissue Simulations – Local Perturbations at Varying Locations

In the previous section we found that a local perturbation with a temperature of T_2_=28°C yielded the optimal performance in terms of rotor attraction quality. Here we model such a perturbation, while we vary its location to study the quality of rotor attraction as a function of the initial distance, *d* [mm], between the perturbation and the core of the spiral wave. The mean tip trajectories during 15 seconds of activity are shown in Fig. 8A for *d* between 5 and 11mm, while the full trajectories as well as their corresponding distance from the perturbation center as a function of time are given in Fig. S3. Rotor attraction was successful only at or below a critical distance of *d*_*cr*_ = 9mm. Over this threshold, no significant effect of the perturbation on rotor meandering or drifting was observed. As expected, when attraction was successful, it was faster as *d* was smaller. The slow transient interval (*STI*) was calculated for each case as in the previous section and plotted against *d* in Fig. 8B. The showed an exponentially decreasing dependence on *d*, following the following best-fitted relationship (R^2^=0.9995),

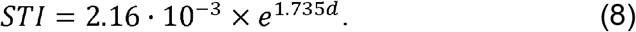

**Fig. 8.**
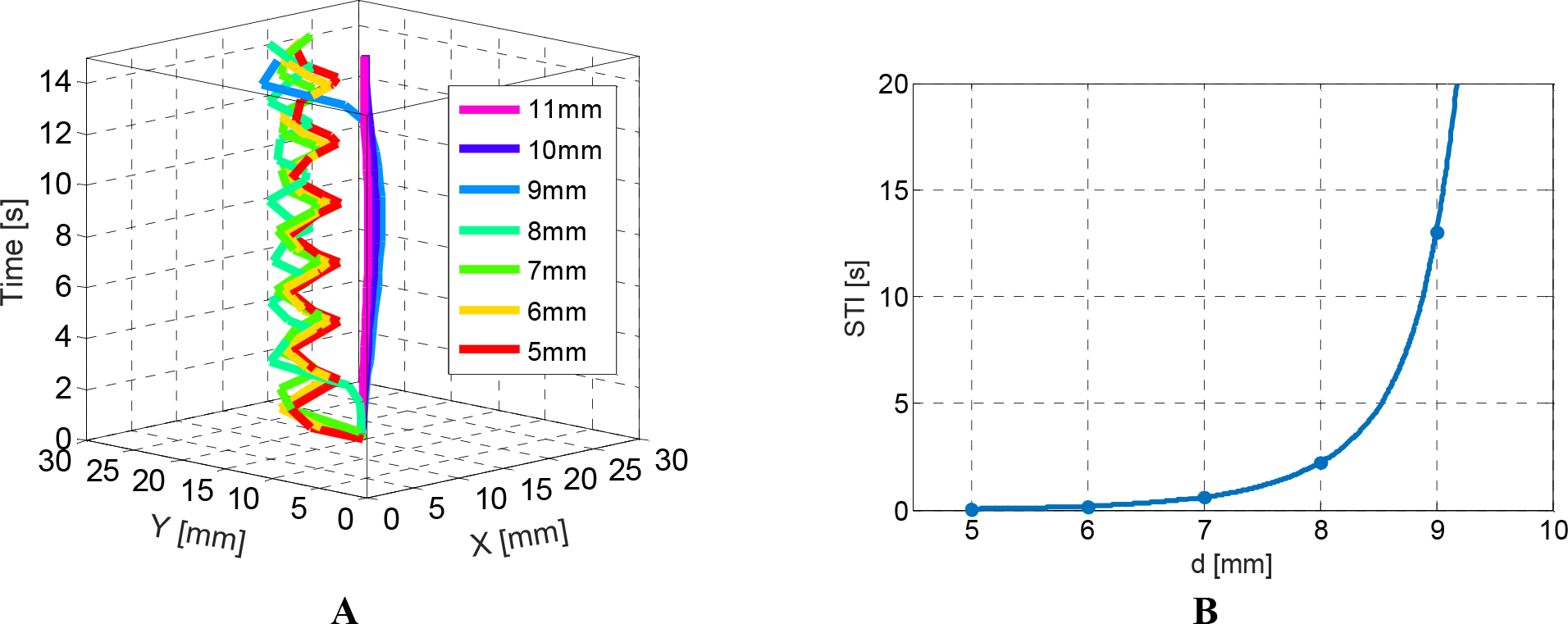
Attraction of a spiral wave using a local perturbation of T2=28°C and varying locations. A. Time-space plots of the tip trajectory. B. Duration of the slow transient interval (STI) phase of attraction as a function of the initial distance, d, between the spiral wave core center and the center of perturbation.

Following this relationship, the attraction of a rotor at an initial distance of d=10mm is expected to require a perturbation application for more than 70sec, which seems to be clinically impractical.

## 4. DISCUSSION

In this study, the effects of externally applied spatial temperature gradients on basic electrophysiological properties of a single atrial myocyte and on spiral wave dynamics in the atrial tissue were numerically studied using a biophysical model of a human atrial tissue. Ion channel remodeling corresponding to chronic AF conditions were employed. Our main findings are as follows: 1) in a single cell with incorporated AF remodeling, temperature exerts a significant effect on recovery time from activation, such that recovery time increases exponentially with decreasing temperatures. In contrast, APD is not affected by temperature in AF conditions; 2) when a linear temperature gradient is applied to the tissue, existing spiral waves drift in a predictable pattern, with one velocity vector component that is parallel and proportional to the negative of the temperature gradient and the other component depends on the rotation direction of the spiral; overall – spirals drift towards the colder region in the tissue, where excitability (as indicated by the sodium channel availability) obtains a global minimum; and 3) a local small temperature perturbation can be employed to attract and anchor a distant spiral wave. The feasibility and quality of such attraction depends on both the temperature of the perturbed tissue as well as on the initial distance between the perturbation and the center of the spiral wave.

Previous experimental and numerical studies have shown that spiral waves can drift in the existence of diffusivity gradient, when the shape of the AP is spatially uniform [42], or in the presence of ionic heterogeneities toward regions with reduced excitability, where an APD gradient is established. Ten Tusscher and Panfilov [27] have shown using a ventricular numerical model that in the presence of a spatial gradient in the APD due to local variations in the K+ currents, spiral waves drifted toward regions of longer spiral wave period. In accordance with our study, the drift velocity consisted of a component along the gradient that was proportional to its magnitude, and a perpendicular component with a direction that depended on the direction of the spiral wave rotation. In line with these results, Calvo et. al. [32] demonstrated that spatial heterogeneity of IK1 channel distribution may cause rotors to drift in the pulmonary vein – left atrial junction (PV-LAJ). This effect was attributed to a gradient in excitability near the rotor pivoting point. Still, both these studies as well as other studies [30-31] were motivated by inherent electrophysiological gradients in the cardiac tissue that may potentially initiate or alter the dynamics of arrhythmias. To the best of our knowledge, the idea to exploit the notion that rotors drift towards low excitability regions in order to artificially attract and anchor rotors for potential ablation applications is novel. However, artificially inducing gradients in the ion channel densities is not feasible clinically. In contrast, imposing local temperature perturbation to the tissue via an inserted catheter is simple. Since temperature sustains a significant effect on the rate constant of biological reactions, it can be used to modulate cellular electrophysiological properties. Indeed, our results show that temperature variations had a substantial effect on the cellular recovery time in both normal and chronic AF remodeled cells (Fig. 2). Noteworthy, while temperature also had a significant effect on the APD in normal cells, such effect was negligible for the AF remodeled cells. Nevertheless, we found that the large sensitivity of the recovery time due to temperature variations was sufficient to induce strong enough spatial gradients of excitability and induce spiral wave drifting in the 2D models.

Some experimental support to our results, indicating that rotor drifting can be induced by a local temperature gradient, can be found in a recent study by Yamazaki et al. [26]. In their experiments, conducted on the left ventricle of the rabbit heart, rotors that were underlying sustained ventricular tachycardia were transformed by regional cooling from stationary to nonstationary when regional cooling was applied. The temperature in the target area was ~30.1°C±1.3°C using a large cooling device with a diameter of 10mm. The authors found that the regional cooling created local long refractoriness due to the prolongation of the APD and the reduction of conduction velocity. Thus, cooling was demonstrated to be capable of unpinning the rotors, drift along the periphery of the regional cooling region, and consequently facilitate termination of reentrant activity by collisions with the boundaries. Our simulations using local temperature perturbations generally support these findings (Fig. 6), however without the rotor termination, apparently since the RTP parameters in the simulations were not similar to those of the experiments. Therefore, we tried to reproduce the experiment protocol by applying RTP with T2=30°C and d=9mm (which is the most distant location that can cause a drift for this type of RTP, as have been found in the study). By applying these RTP protocol we succeeded to imitate the same behavior, so the rotor had been destabilized, drifted toward the colder region, and eventually terminated as a consequence of collision with the upper border (Fig. 9). Also, in contrast to the experiments, we show that spiral wave drift is obtained even in the AF remodeled tissue, where temperature variation sustains no significant effect on the APD but only on the recovery time. Also, our model revealed that the tip trajectory velocity was slower as the temperature decreased, and its rotation period per cycle increased (Fig. S1). These findings, along with the net drift demonstration in Fig. 5, are correlate to the notion that lower temperature regions yielding a gradient in the tissue excitability, and therefore slow the rotor motion and require a longer drifting distance before the excitability level permits the rotor to reenter back. We further found that an optimal effect in terms of spiral wave attraction time (STI, slow transient interval) and the mean distance of the anchored spiral to the perturbation center (MD, mean distance) can be obtained for a temperature of 28°C (Fig. 7). While MD decreased monotonically with decreasing perturbation temperatures (Fig. 7A), the STI was non-monotonous, and obtained a minimum at 28°C, thus defining the optimal perturbation temperature. As we showed in Fig. 3 for linear temperature gradients, drifting velocity was proportional to the spatial temperature gradient. Moreover, in accordance with our hypothesis, the underlying ionic mechanism was linked to the spatial gradient in the tissue excitability due to the temperature gradient (Fig. 4). To mechanistically understand our observation that a local 28°C temperature perturbation provides the optimal, shortest drifting time (or STI) we plotted in Fig. 10A the time-averaged sodium channel availability (mean *h*×*j*, averaged during a single spiral wave rotation) along the red dashed line marked in Fig. 6B (left) for all the simulations described in Fig. 6. This line connects between the initial center of the spiral wave and the center of perturbation and thus represents an optimal drifting path for the spiral wave. As Fig. 10A shows, the mean *h*×*j* profile is characterized by a local trough at y~15mm that corresponds to the low-excitability spiral wave core center, and a region of low *h*×*j* values between y=22mm and y=24mm that corresponds to the location of the perturbation. The dashed black lines in Fig. 10A mark the region between y=17mm - the right peak to the right of the trough (or the edge of the spiral wave core) - and y=22mm - the left edge of the perturbation. That region contains the spatial gradient of *h*×*j* that is affecting the spiral wave drifting. For each temperature, the mean *h*×*j* gradient in that region was estimated by the slope of the best fit line between those two dashed lines. As shown in Fig. 10B, the drifting time (STI) was proportional to the negative of the *h*×*j* gradient. A linear regression analysis (Fig. 10C) showed indeed that these two properties were linearly correlated with the following relationship (R^2^=0.83),

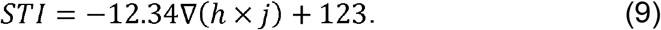

Thus, for gradually decreasing perturbation temperatures the gradient of excitability as sensed by the spiral wave initially slowly increased, reaching a plateau at T2=28°C. Below that temperature, the gradient magnitude decreased rapidly. As a result, the drifting time, represented by the STI, which is correlated to the excitability gradient by (9), rapidly increased, deteriorating the quality of attraction.

**Fig. 9.**
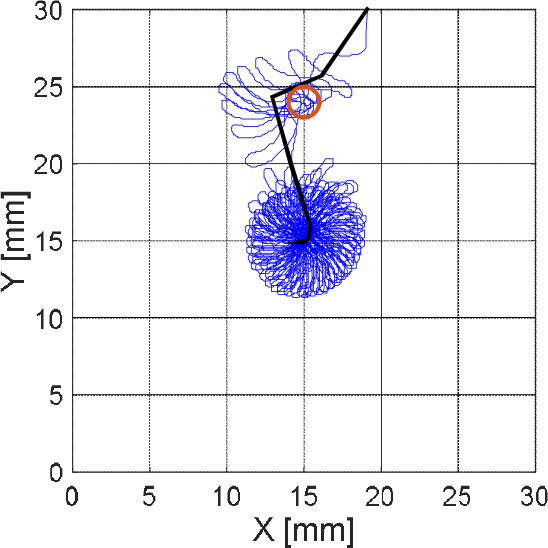
Employing RTP with T_2_=30°C and d=9mm, yielding the rotor to be destabilized, drift toward the colder region, and eventually terminated as a consequence of collision with the upper border (at Y=30mm).

**Fig. 10.**
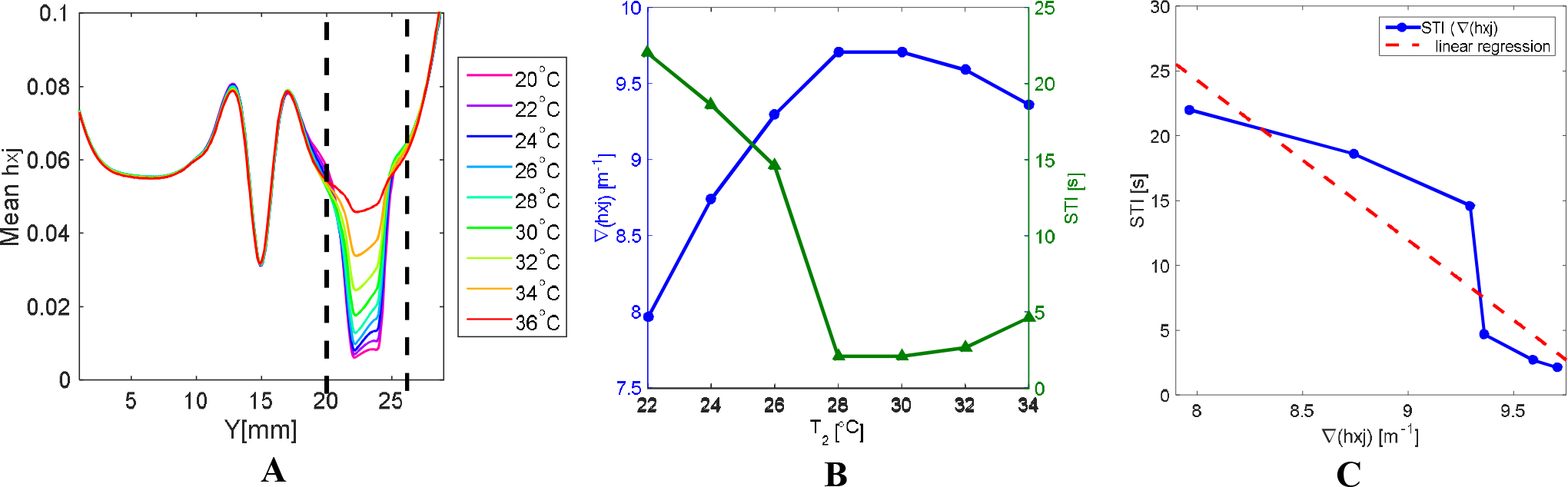
Analysis of spiral wave drifting mechanism for a local temperature perturbation. A. Mean sodium channel availability along the dashed vertical line in Fig. 6B (left). The region between the two dashed black lines refers to the vertical distance between the spiral wave center and the perturbation center, and the gradient of excitability in that region was found to correlate with the drifting time of the spiral wave before anchoring to the perturbation. B. The gradient of excitability (between the two dashed lines in panel A) and the STI as a function of perturbation temperature. C. Relationship between the gradient of excitability and the STI reveals a decreasing monotonic function.

### Limitations

In this work, the CRN model was employed with AF modifications in an isotropic, electrophysiologically uniform 2D tissue to yield a single rotor activity. Other AF models, e.g., the model employed by Jacquemet et al [43], integrated anatomically and dynamically complex elements like 3D geometry, tissue anisotropy and other physiological heterogeneities that yielded complex AF conduction patterns such as multiple wavelets, spiral wave breakups and collisions. Yet, the aim of our study was to establish the most fundamental theory underlying the relationship between temperature gradients and rotor drifting. Thus, it was in fact our intention to employ a simplified, isotropic anatomy model wherein a single rotor is active in order to allow a direct analysis of such relationships, while avoiding interfering factors that will render such analysis too complex. In future studies we will utilize the mechanisms established in this paper to further analyze the effects of temperature heterogeneities in a more complex anatomy and AF electrical propagation patterns. It should also be noted that while a single rotor is indeed a relatively simple kind of propagation dynamics during AF, the “mother rotor” hypothesis is one of the potential suggested mechanism sustaining AF, that supported by the exhibition in the AF driving region of maximum dominant frequency [44]. Therefore, simulating persistent AF driver as a single rotor may not be a simplified scenario, but rather a likely one.

### Potential clinical implications and future studies

Our study aims at understanding the basic mechanisms of spiral wave drifting and attraction in the presence of applied temperature perturbations, and the presented results cannot be directly translated into the clinical setup. Still, this study was motivated by the potential prospective clinical application of the proposed method, and it is our hope that the mechanistic insights from our study may be utilized in the future as a proof-of-concept in the design of a new methodology for AF characterization and termination. Surely such application should be comprehensively investigated in future experimental and clinical studies. We believe that the interaction between rotors and temperature gradients can be utilized to help tracking rotors. For example, it is well-known that rotor drifting leading to Doppler-induced differences in local activation periods along the direction of drift [45]. Therefore, it is hypothesized that the application of local temperature perturbation while measuring the electrical activity at several few pre-set locations may allow to back-track the origin of a drifting rotor. This can be theoretically achieved by measuring the Doppler shifts at the various locations, which arise only due to the rotor drifting rather than stable rotor or ectopic source, and applying an inverse reconstruction algorithm. However, causing for a controlled spiral wave drift in the clinic is on the verge of impracticality, and the method presented in this study can provide a novel solution for it. The development of such an algorithm, in different scenarios, is the goal of our future research. A probable clinical scenario would be the application of temperature gradients in a fibrotic atrial tissue. As rotors tend to stabilize around areas of fibrotic tissue or tissue heterogeneities, a question arises whether and under what configurations, reentrant waves can or cannot detach from fibrotic patches due to applied temperature perturbations. Such a question should be undoubtedly investigated in the future. Finally, future research should have to investigate if application of local cooling in persistent AF patients and using the Doppler Effect will enable to characterize the type of driving arrhythmogenic source, being ectopic foci (and thus not being affected by the applied temperature gradients and therefore not drifting) or a rotor that is expected to drift in the presence of temperature gradient. We predict that a controlled spiral wave drift will occur under spatial temperature gradient, causing to Doppler Effect that can be used for analyze the arrhythmogenic source type. This interesting classification problem should be also studied in order to fully understand the limitations and potential of local temperature application in the characterization of AF activity.

Competing interests: None declared

Funding: None

Ethical approval: Not required

---

**Figure S1.**
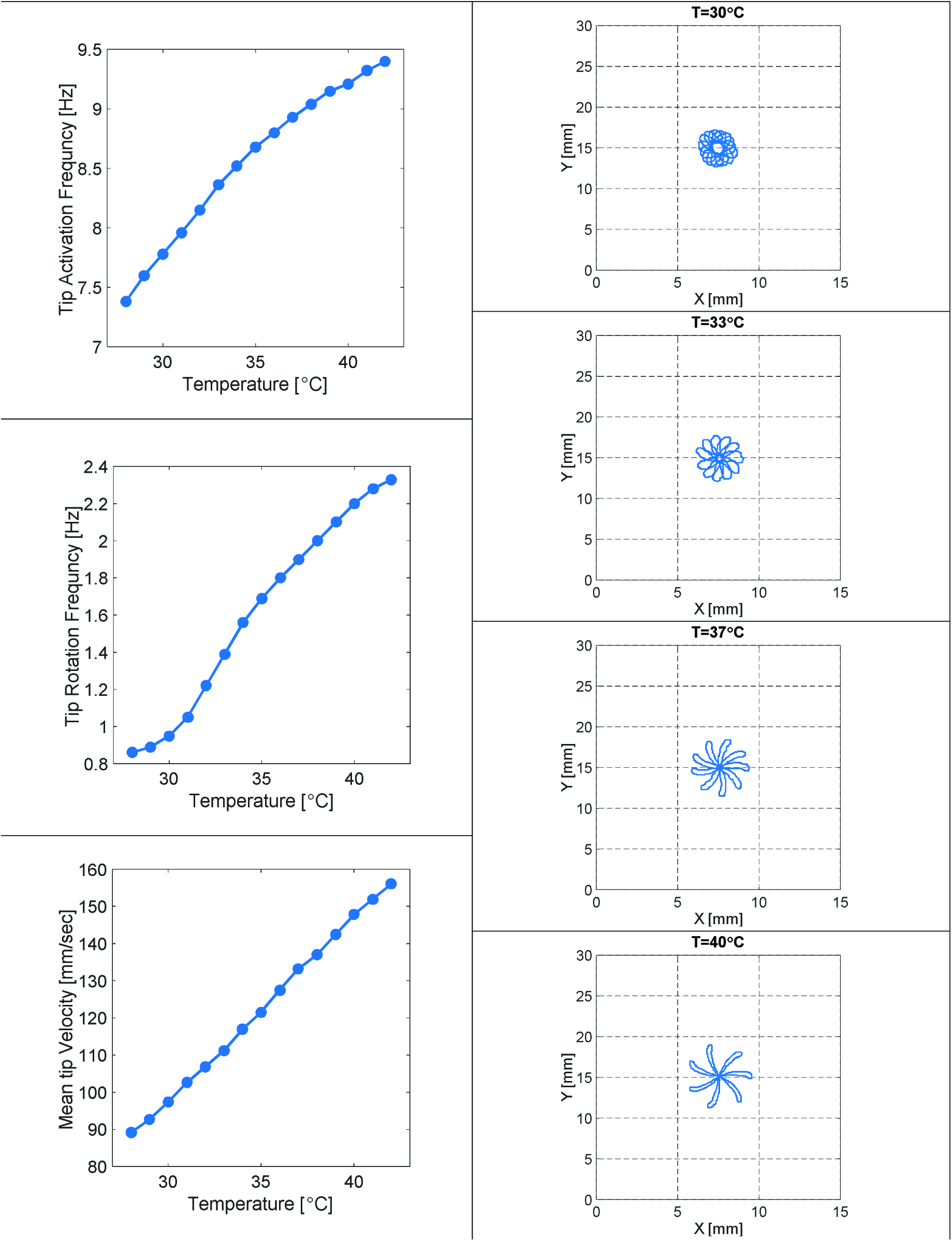
Spiral wave properties as function of constant tissue temperature. Left column: Rotor activation frequency (f1), which is the fast frequency that determines the interval between sequential local excitation (top), spiral wave tip rotation frequency (f2) (middle), which is the slow frequency that determines the rotation rate of a rotor, (middle), and the mean tip velocity (bottom). Spiral wave dynamics exhibited a monotonic behavior as function of temperature, so that the wavefront propagation is being slower and have smaller frequencies with decreasing temperatures. Right column: Spiral wave tip trajectory of one rotation cycle for several cases (T equal to 30°C, 33°C, 37°C and 40°C), demonstrated the different trajectory patterns as function of temperature.

**Figure S2.**
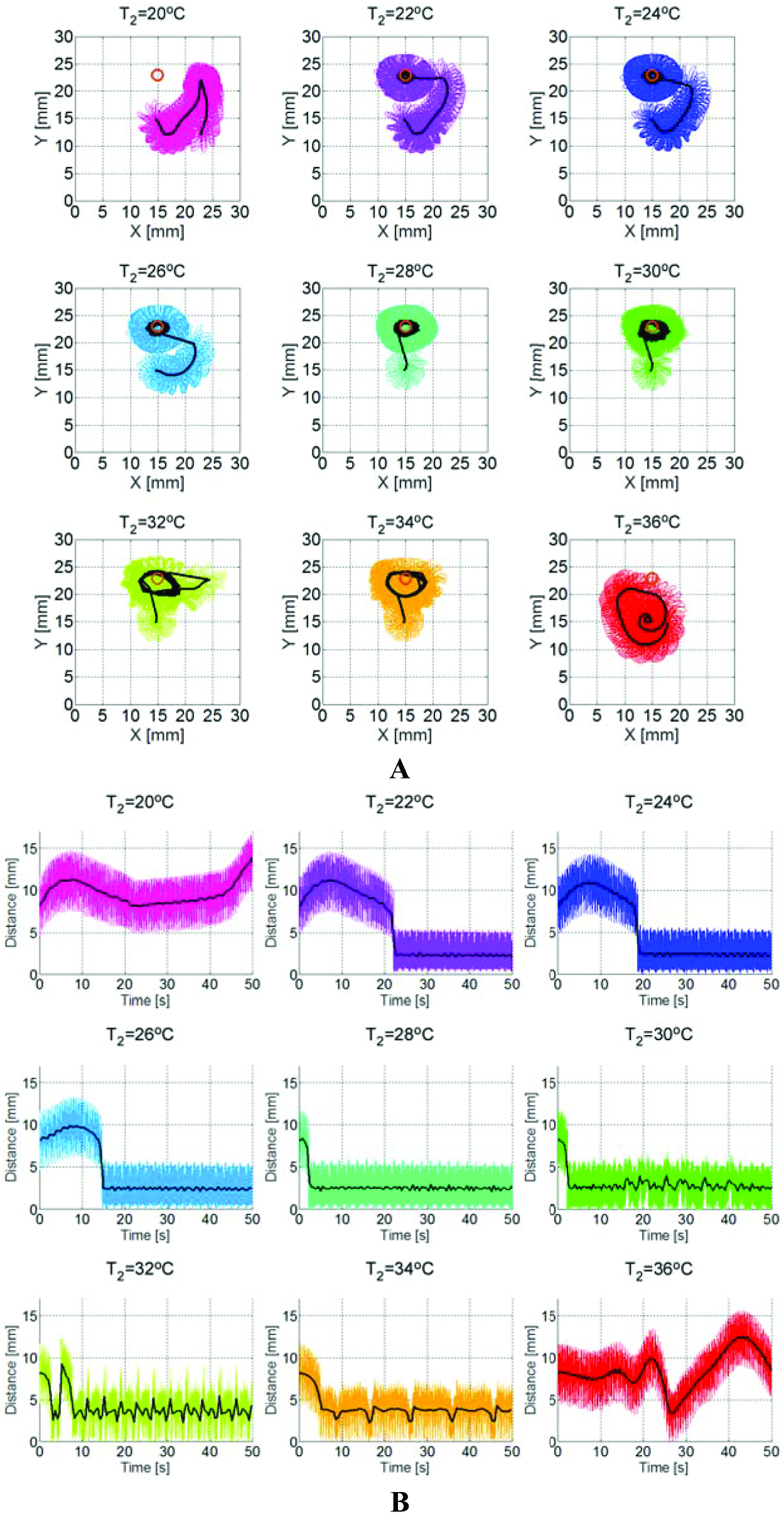
Simulations of local perturbations of varying temperatures. A. Tip trajectories of a spiral wave in the presence of the local perturbations. In black – mean trajectories. Perturbation location is marked by a circle. B. Distance of the mean tip trajectory from the center of perturbation as a function of time.

**Figure S3.**
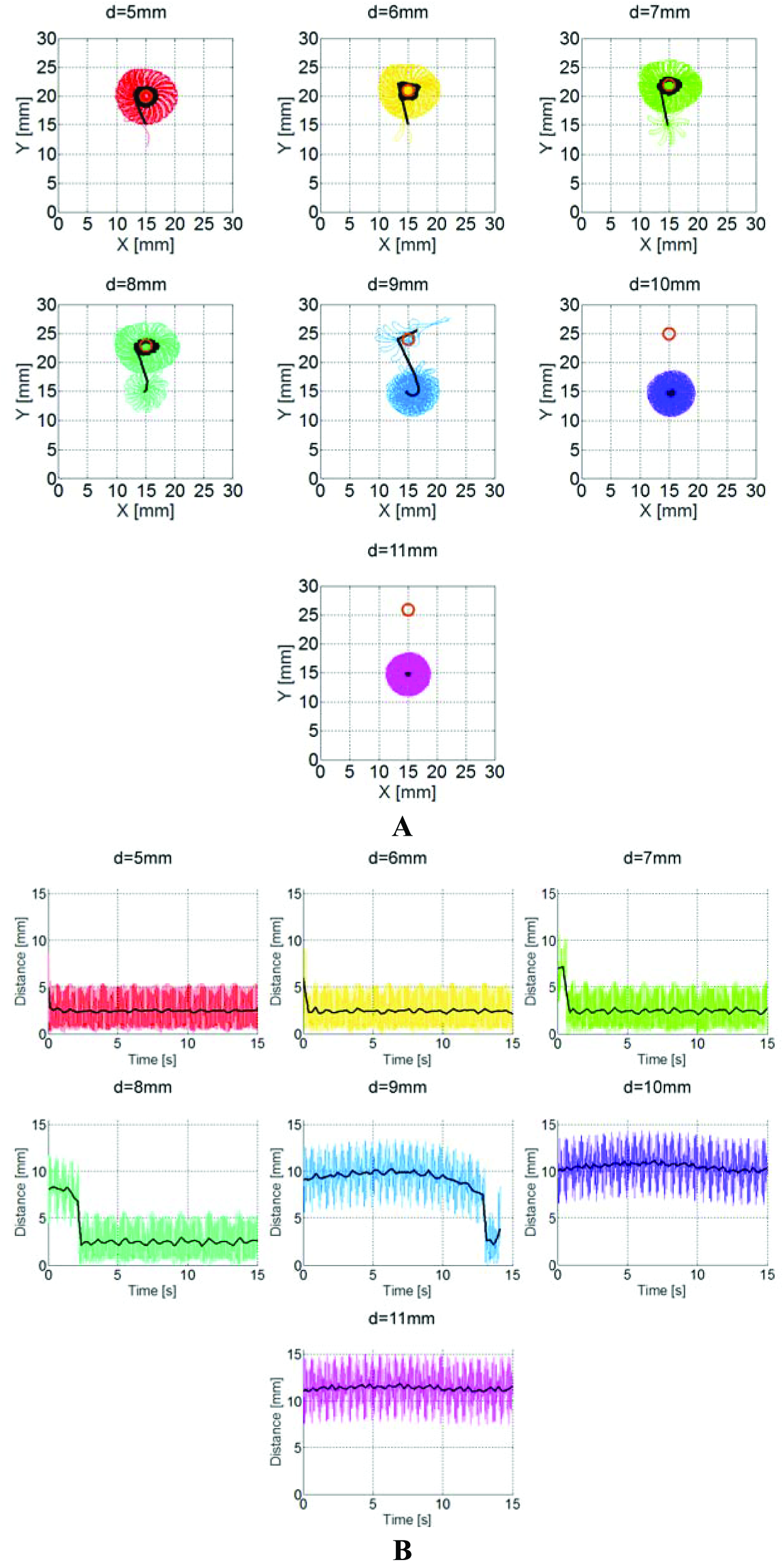
Simulations of local perturbations of varying initial distance from the spiral wave core. A. Tip trajectories of a spiral wave in the presence of the local perturbations. In black – mean trajectories. Perturbation location is marked by a circle. B. Distance of the mean tip trajectory from the center of perturbation as a function of time.

